# Provitamin A maize inbred lines exhibit resistance to multiple foliar diseases under laboratory and field conditions

**DOI:** 10.64898/2026.09.03.749191

**Authors:** Mamadou Mboup, Adefoyeke O. Aduramigba-Modupe, Bunmi Olasanmi, Wende Mengesha, Silvestro Meseka, Ibnou Dieng, Abebe Menkir, Alejandro Ortega-Beltran

## Abstract

Maize (*Zea mays*) is a staple food for millions in sub-Saharan Africa (SSA), contributing to both caloric intake and essential micronutrients. Recent breeding efforts have focused on enhancing its nutritional value by increasing provitamin A (PVA) carotenoid content to address vitamin A deficiency (VAD), a common problem among children under 5 years, pregnant and lactating mothers across SSA. However, high rainfall, and both warm and humid conditions in various SSA regions result in devastating foliar diseases such as maize streak virus (MSV), northern corn leaf blight (NCLB), southern corn leaf blight (SCLB), southern corn rust, grey leaf spot (GLS), and Curvularia leaf spot (CLS). Breeding for resistance can aid in mitigating yield losses caused by those diseases. Rapid, efficient screening methods can allow for examining large germplasm collections. The current study evaluated 21 maize inbred lines with contrasting PVA content, along with two commercial inbred controls, for resistance to *Exserohilum turcicum*, *Bipolaris maydis*, and *Curvularia lunata*, causal agents of NCLB, SCLB, and CLS, respectively, using a detached leaf assay (DLA) and under natural field infestation. Significant variation was detected among the inbreds, seven were classified as resistant to certain diseases, ten as moderately resistant, and six as susceptible to all foliar diseases. Overall, high PVA inbred lines had less susceptibility to NCLB, SCLB, and CLS. The results suggest that PVA-enriched maize inbred lines possess improved resistance to multiple foliar diseases. The field assessments of disease severity validated the effectiveness of the DLA in distinguishing resistant from susceptible inbred lines, and resistance to other diseases in field conditions was detected in parallel. The results indicate that high PVA maize has the potential to simultaneously address VAD, mycotoxin contamination in SSA with improved resistance to multiple foliar diseases.

## Introduction

Maize (*Zea mays* L.) is a leading staple cereal crop cultivated worldwide providing about a quarter of the total caloric intake and essential micronutrients to over 4.5 billion people (FAO, 2022; Shiferaw et al., 2011; WHO, 2009). The crop constitutes a fundamental part of daily diets in developing countries, particularly in sub-Saharan Africa (SSA). Maize is projected to continue increasing in area under production, with an annual rate of 1.3%, surpassing wheat (0.9%) and other coarse grains (0.6%) (FAO, 2022). In SSA, total maize production is forecasted to reach 24 million metric tons (mmt) in 2025, representing an increase of up to 2.6 mmt from the previous year (FAO, 2022; USDA, 2025).

Given its crucial role in the diet of billions and its increasing importance in livestock feed and biofuel production, breeding programs have prioritized the development of high-yielding biofortified varieties i) aiming to mitigate micronutrient deficiencies [Zn, Fe, Se, I, and provitamin A (PVA)], and ii) supply raw materials to feed and brewery industries. In the case of PVA, breeders have developed and released high PVA maize varieties to alleviate vitamin A deficiency (VAD) (Bouis & Saltzman, 2017; HarvestPlus, 2014; Xue et al., 2023). However, maize is susceptible to various foliar diseases impacting yield and quality in most tropical regions of SSA. Foliar diseases can cause extensive leaf damage, including chlorotic streaks, necrotic lesions and secondary infections, which reduce photosynthesis, accelerate defoliation and senescence, and may lead to significant yield losses ranging from 15% to 100% (Badu-Apraku & Fakorede, 2017; Akinwale & Oyelakin, 2018). Among the most devastating maize foliar diseases in SSA are northern corn leaf blight (NCLB), southern corn rust (SCR), southern corn leaf blight (SCLB), maize streak virus (MSV), grey leaf spot (GLS), and Curvularia leaf spot (CLS) (Wang et al., 2014; Badu-Apraku & Fakorede, 2017).

NCLB, caused by *Exserohilum turcicum*, occurs across SSA’s mid- and high-altitude regions, and more recently, in lowland zones of West and Central Africa. The pathogen thrives under high humidity (75%–90%) and moderate temperatures (22°C–25°C) (Ahangar et al., 2022) producing necrotic spindle-shaped spots (Badu-Apraku & Fakorede, 2017; Akinwale & Oyelakin, 2018). NCLB early infection can reduce maize yields by >90% (Ahangar et al., 2022). SCLB, induced by *Bipolaris maydis*, is another highly destructive disease characterized by diamond-shaped lesions that may coalesce to blight entire leaves under warm, humid conditions. Although its prevalence is relatively limited across SSA, it poses a threat in southwestern Nigeria and has historically caused major losses, including a 15% drop in national maize yield in the USA (Ullstrup, 1972; Singh et al., 2012; Akinwale & Oyelakin, 2018). CLS, caused by *Curvularia lunata*, is prevalent in tropical high-rainfall regions, inducing scattered necrotic or chlorotic spots that may cover up to 60% of the leaf surface, thereby reducing photosynthetic area (Wang et al., 2014).

Other important diseases include MSV, vectored by *Cicadulina* spp., which is endemic in SSA and capable of causing up to 100% yield loss in susceptible varieties (Emeraghi et al., 2021). GLS, caused by the necrotrophic *Cercospora zeae-maydis*, is a polycyclic disease that can occur under cool, humid conditions and has caused regional outbreaks of up to 30% yield loss (Diro & Lemessa, 2022). SCR is caused by Puccinia polysora, a destructive biotrophic pathogen prevalent in hot, humid tropical and subtropical lowlands, where it promotes premature senescence, leading to yield losses exceeding 50% under optimal infection conditions (Rhind et al., 1952; Badu-Apraku & Fakorede, 2017; Cao et al., 2024). *Colletotrichum graminicola* is a global distributed fungus causing anthracnose leaf blight (ALB), stalk rot and top dieback, particularly in warm, humid environments. ALB can occur throughout maize development and has been reported to reduce grain yield by up to 20% in hybrids and 30% in inbreds (Badu-Apraku et al., 1987; CIMMYT, 2004).

While maize diseases can be mitigated through the application of a range of fungicides and biocontrol agents, host plant resistance remains the most cost-effective and sustainable approach for controlling diseases, especially for maize cultivated by resource-constrained smallholder farmers. Accordingly, maize breeding programs have prioritized routine germplasm screening, resulting in breeding materials with high to moderate levels of resistance to several diseases (Badu-Apraku & Fakorede, 2017). However, despite these efforts, devastating disease outbreaks still occur unpredictably due to pathogens’ mutations, the emergence of new races of pathogens, or the introduction of new ecotypes, trends increasingly influenced by climate change.

To strengthen maize resilience, especially in biofortified germplasm, there is a need to sufficiently characterize resistance in PVA inbred lines against multiple foliar diseases. Detached leaf assay (DLA) has been reported as a reliable, rapid, and effective approach for rapidly screening a large set of genotypes under controlled conditions while ensuring uniform disease development with standardized inoculum loads. DLA also allows for the simultaneous testing of multiple isolates on the same genotype (Degani & Cernica, 2014; Alakonya et al., 2018; Aregbesola et al., 2020; Bankole et al., 2022). However, resistance must be confirmed under field conditions in endemic hotspots where natural infection pressure confirms practical resistance performance. Therefore, the objectives of the current study were to i) rapidly assess in DLA various PVA inbred lines for resistance to NCLB, SCLB, CLS, and ii) evaluate their field resistance to multiple diseases (NCLB, SCLB, CLS, MSV, GLS, SCR, and ALB) under natural conditions. The obtained results will support the development of high PVA maize with higher levels of durable multiple disease resistance to strengthen productivity and food safety and security in SSA.

## Materials and methods

### Plant materials

Twenty-three maize inbred lines with PVA content ranging from 5.4 to 51.7 µg/g developed by the Maize Improvement Program at IITA (MIP-IITA) were evaluated for resistance to multiple foliar diseases. Ten of the inbred lines possess desirable general combining ability (GCA) for β-carotene and PVA, and resistance to both *A. flavus* infection and aflatoxin production, while nine inbred lines have undesirable, inconsistent GCA for these traits (Mboup et al., 2023). Also, the nine high PVA inbred lines accumulated low fumonisin in field trials (Mboup et al., 2024). Two released aflatoxin resistant inbred lines, tolerant to multiple foliar diseases, along with two inbred line testers with contrasting PVA content (low = 14.4 μg/g; high = 25.0 μg/g) used to discriminate PVA inbred lines for PVA content and aflatoxin accumulation, were included as checks in the experiments (Table 1). The inbred lines were divided into two groups: high (≥25 µg/g) and low (< 25 µg/g) PVA, hereafter referred to as H and L, respectively.

**Table 1.**
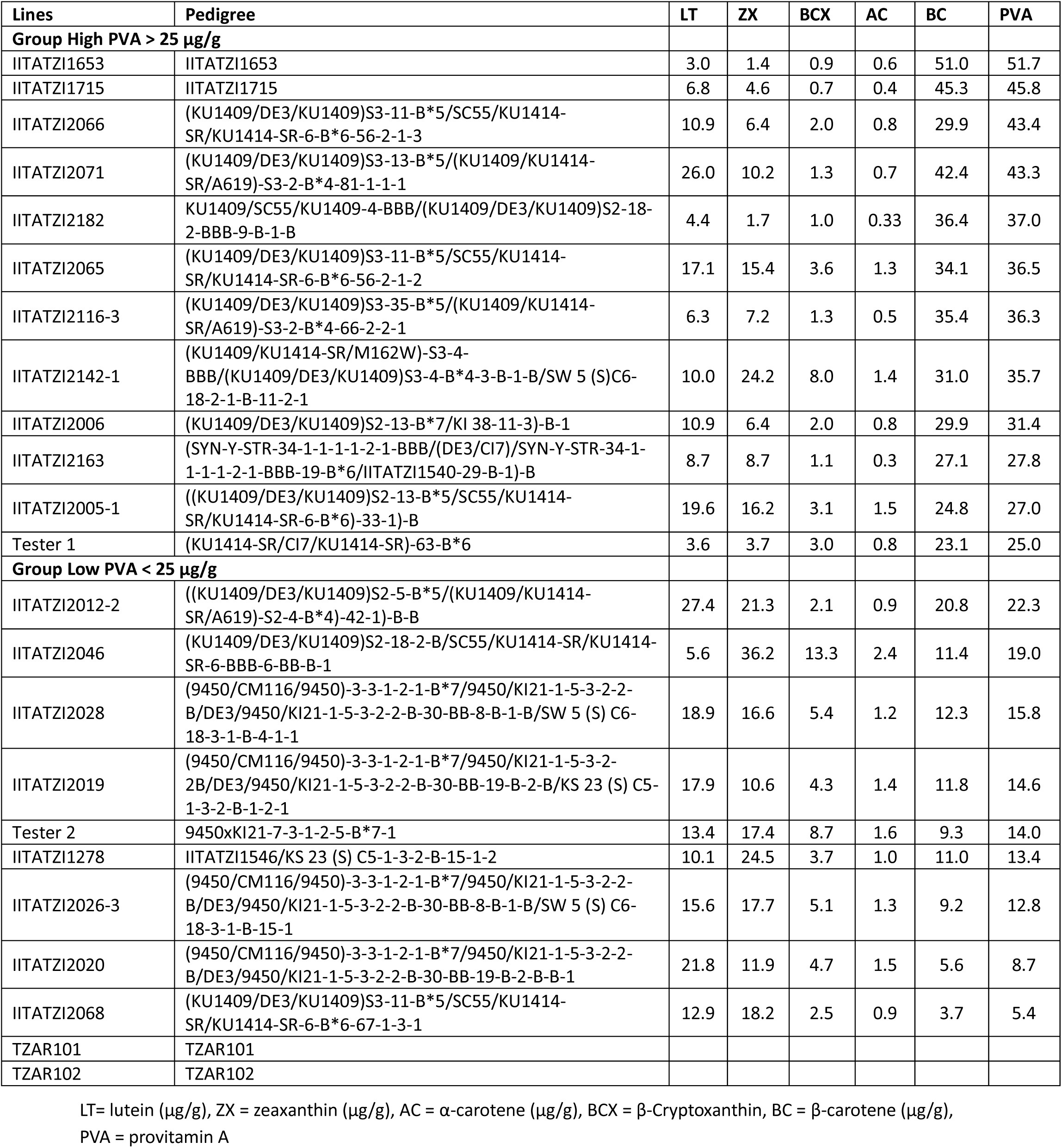
PVA maize inbred lines selected for evaluation of resistance to multiple foliar diseases using a detached leaf assay and under natural field conditions.

| Lines | Pedigree | LT | ZX | BCX | AC | BC | PVA |
| --- | --- | --- | --- | --- | --- | --- | --- |
| **Group High PVA > 25 µg/g** |  |  |  |  |  |  |  |
| IITATZI1653 | IITATZI1653 | 3.0 | 1.4 | 0.9 | 0.6 | 51.0 | 51.7 |
| IITATZI1715 | IITATZI1715 | 6.8 | 4.6 | 0.7 | 0.4 | 45.3 | 45.8 |
| IITATZI2066 | (KU1409/DE3/KU1409)S3-11-B*5/SC55/KU1414-SR/KU1414-SR-6-B*6-56-2-1-3 | 10.9 | 6.4 | 2.0 | 0.8 | 29.9 | 43.4 |
| IITATZI2071 | (KU1409/DE3/KU1409)S3-13-B*5/(KU1409/KU1414-SR/A619)-S3-2-B*4-81-1-1-1 | 26.0 | 10.2 | 1.3 | 0.7 | 42.4 | 43.3 |
| IITATZI2182 | KU1409/SC55/KU1409-4-BBB/(KU1409/DE3/KU1409)S2-18-2-BBB-9-B-1-B | 4.4 | 1.7 | 1.0 | 0.33 | 36.4 | 37.0 |
| IITATZI2065 | (KU1409/DE3/KU1409)S3-11-B*5/SC55/KU1414-SR/KU1414-SR-6-B*6-56-2-1-2 | 17.1 | 15.4 | 3.6 | 1.3 | 34.1 | 36.5 |
| IITATZI2116-3 | (KU1409/DE3/KU1409)S3-35-B*5/(KU1409/KU1414-SR/A619)-S3-2-B*4-66-2-2-1 | 6.3 | 7.2 | 1.3 | 0.5 | 35.4 | 36.3 |
| IITATZI2142-1 | (KU1409/KU1414-SR/M162W)-S3-4-BBB/(KU1409/DE3/KU1409)S3-4-B*4-3-B-1-B/SW 5 (S)C6-18-2-1-B-11-2-1 | 10.0 | 24.2 | 8.0 | 1.4 | 31.0 | 35.7 |
| IITATZI2006 | (KU1409/DE3/KU1409)S2-13-B*7/KI 38-11-3)-B-1 | 10.9 | 6.4 | 2.0 | 0.8 | 29.9 | 31.4 |
| IITATZI2163 | (SYN-Y-STR-34-1-1-1-2-1-BBB/(DE3/CI7)/SYN-Y-STR-34-1-1-1-2-1-BBB-19-B*6/IITATZI1540-29-B-1)-B | 8.7 | 8.7 | 1.1 | 0.3 | 27.1 | 27.8 |
| IITATZI2005-1 | ((KU1409/DE3/KU1409)S2-13-B*5/SC55/KU1414-SR/KU1414-SR-6-B*6)-33-1)-B | 19.6 | 16.2 | 3.1 | 1.5 | 24.8 | 27.0 |
| Tester 1 | (KU1414-SR/CI7/KU1414-SR)-63-B*6 | 3.6 | 3.7 | 3.0 | 0.8 | 23.1 | 25.0 |
| **Group Low PVA < 25 µg/g** |  |  |  |  |  |  |  |
| IITATZI2012-2 | ((KU1409/DE3/KU1409)S2-5-B*5/(KU1409/KU1414-SR/A619)-S2-4-B*4)-42-1)-B-B | 27.4 | 21.3 | 2.1 | 0.9 | 20.8 | 22.3 |
| IITATZI2046 | (KU1409/DE3/KU1409)S2-18-2-B/SC55/KU1414-SR/KU1414-SR-6-BBB-6-BB-B-1 | 5.6 | 36.2 | 13.3 | 2.4 | 11.4 | 19.0 |
| IITATZI2028 | (9450/CM116/9450)-3-3-1-2-1-B*7/9450/KI21-1-5-3-2-2-B/DE3/9450/KI21-1-5-3-2-2-B-30-BB-8-B-1-B/SW 5 (S) C6-18-3-1-B-4-1-1 | 18.9 | 16.6 | 5.4 | 1.2 | 12.3 | 15.8 |
| IITATZI2019 | (9450/CM116/9450)-3-3-1-2-1-B*7/9450/KI21-1-5-3-2-2-B/DE3/9450/KI21-1-5-3-2-2-B-30-BB-19-B-2-B/KS 23 (S) C5-1-3-2-B-1-2-1 | 17.9 | 10.6 | 4.3 | 1.4 | 11.8 | 14.6 |
| Tester 2 | 9450xKI21-7-3-1-2-5-B*7-1 | 13.4 | 17.4 | 8.7 | 1.6 | 9.3 | 14.0 |
| IITATZI1278 | IITATZI1546/KS 23 (S) C5-1-3-2-B-15-1-2 | 10.1 | 24.5 | 3.7 | 1.0 | 11.0 | 13.4 |
| IITATZI2026-3 | (9450/CM116/9450)-3-3-1-2-1-B*7/9450/KI21-1-5-3-2-2-B/DE3/9450/KI21-1-5-3-2-2-B-30-BB-8-B-1-B/SW 5 (S) C6-18-3-1-B-15-1 | 15.6 | 17.7 | 5.1 | 1.3 | 9.2 | 12.8 |
| IITATZI2020 | (9450/CM116/9450)-3-3-1-2-1-B*7/9450/KI21-1-5-3-2-2-B/DE3/9450/KI21-1-5-3-2-2-B-30-BB-19-B-2-B-B-1 | 21.8 | 11.9 | 4.7 | 1.5 | 5.6 | 8.7 |
| IITATZI2068 | (KU1409/DE3/KU1409)S3-11-B*5/SC55/KU1414-SR/KU1414-SR-6-B*6-67-1-3-1 | 12.9 | 18.2 | 2.5 | 0.9 | 3.7 | 5.4 |
| TZAR101 | TZAR101 |  |  |  |  |  |  |
| TZAR102 | TZAR102 |  |  |  |  |  |  |

LT= lutein (µg/g), ZX = zeaxanthin (µg/g), AC = α-carotene (µg/g), BCX = β-Cryptoxanthin, BC = β-carotene (µg/g),
PVA = provitamin A

### Fungi and inoculum preparation

The foliar pathogens used for DLA (see below) were *E. turcicum*, *B. maydis*, and *C. lunata*, the causal agents of NCLB, SCLB, and CLS, respectively. Two isolates each of *C. lunata* (CV1, CV2) and *E. turcicum* (NGETIB16-13, NGETIK16-12), and one isolate of *B. maydis* (SLB-05) were used to inoculate sections of detached leaves of each PVA inbred line and the control genotypes. These isolates are part of the fungal collection of IITA Pathology and Mycotoxin Unit and their virulence and utility in DLA have been previously reported (Aregbesola et al., 2020; Bankole et al., 2022).

All the isolates were independently grown on media containing 20% V8 Juice®, 1.5 g/l CaCO_3_, and 20 g/l agar, and incubated for 14 days. Then, conidia were washed with a sterile TWEEN®20 solution (2-3 drops/100 ml sterile distilled water), filtered, and adjusted to a 10^5^ conidia/ml concentration for each isolate using a hemacytometer.

### Agar medium preparation for DLA

Amended Technical Agar (TA) media was prepared following the methods described in previous studies in our lab (Alakonya et al., 2018; Aregbesola et al., 2020; Bankole et al., 2022). Briefly, Technical Agar (10 g/l) was autoclaved at 121°C for 20 min and thereafter amended aseptically inside a biosafety cabinet with 0.045 g/l of 6-benzylaminopurine (BAP) phytohormone. Lactic acid (1.5 ml/l) was added to prevent bacterial contamination. After cooling, 250 ml of the media was dispensed into the sterile plastic crisper boxes (23 × 31 × 10 cm), which had been disinfected by washing and soaking in a NaOCl (10%) + mancozeb (12.5 mg/l) solution and rinsed with sterile distilled water.

### Maize screenhouse planting and leaf assay inoculation procedure

The 23 PVA maize inbred lines and the control genotypes were assessed for resistance to multiple foliar diseases using the DLA. All maize materials were planted in a screenhouse in 8-kg pots. Four weeks after planting, fully expanded, healthy leaves of each material were excised and transferred to the laboratory. After rinsing with tap water, the leaves were cut into 14 × 5 cm pieces. The leaves were surface-sterilized in a biosafety cabinet by successive dipping into sterile solutions of 1% NaOCl and 70% NaOH for 90 sec each, followed by rinsing with three changes of sterile distilled water. Thereafter, the excised leaves were blotted with sterile paper towels before being placed in the sterile crisper boxes containing the amended TA media. The adaxial side of the leaves rested on the media.

Each isolate was independently inoculated by applying 40 μl spore suspension (10^5^ conidia/ml) onto two locations on the abaxial side of the excised leaves of the inbred lines. A single isolate was used to inoculate each leaf. Sterile distilled water was used for negative control treatments. Crisper boxes containing the leaves were sealed with cling film and incubated on benchtops at room temperature (25°C) with a 12 hr photoperiod for 10 days. The experiment followed a 3 × 23 factorial design, where the three treatments (two isolates and a mock-inoculated control) were evaluated in the 23 PVA inbred lines, except for *B. maydis*, which had only one isolate. The inoculated leaves were arranged in a completely randomized design (CRD) with each treatment replicated three times. Each disease was evaluated independently using different leaves of the same maize plants, and the experiment was repeated twice.

### Disease assessment and genotype ranking

After inoculation, disease severity (DS) for each foliar pathogen was determined every two days over a 10-day period using a 1 to 5 scale based on the leaf area infected at the point of inoculation. The percentage area affected was visually estimated as follows: 1 = 0–5%; 2 = >5–10%; 3 = >10– 25%; 4 = >25–50%; and 5 = >50% (Bankole et al., 2022). The DS was used to calculate the area under the disease progress curve (AUDPC) according to the formula of (Campbell and Madden, 1990).

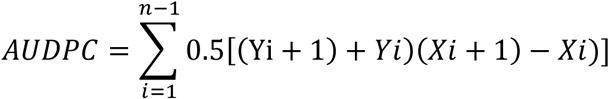

Where Yi = DS (per unit) at the ith observation,

Xi = time (days) at the ith observation, and n = total number of observations.

The AUDPC values were used to compute a rank summation index (RSI) to assess resistance to multiple diseases. The RSI assigned greater absolute values to genotypes exhibiting superior performance. The assigned values were then summed to determine the rank of each genotype. Based on the RSI values, genotypes were classified as resistant (R) (RSI range = 165–213), moderately resistant (MR) (RSI range = 214–229), and susceptible (S) (RSI range = 230–265).

### Field assessment

A field experiment was conducted during the 2023 main cropping season at two IITA research stations in Nigeria: Ikenne (3°42′ E, 6°54′ N, 60 masl) and Ibadan (3°54′ E, 7°29′ N, 227 masl). Ikenne is a hotspot of foliar diseases. It lies in tropical humid rainforest agro-ecological zone of south-west Nigeria, with high average relative humidity (80–85%) and annual rainfall ranging from 1600 to 1800 mm. Similarly, Ibadan is also foliar disease hotspot; it lies in sub-humid tropical forest agro-ecology bordering forest and savanna ecosystems. The two locations have bimodal rainfall pattern allowing for two cropping cycles in year. Details of the field layout and field management have been described in a study where resistance f of the inbred lines Aspergillus ear rot, Fusarium ear rot, and both aflatoxin contamination and fumonisin contamination was evaluated (Mboup et al., 2024). Briefly, a total of 23 inbred lines were planted using a randomized complete block design and replicated three times. Seeds of each entry were planted manually in a 4-m row with a space of 0.75 m between rows and 0.25 m distance between plants at both locations, which were later thinned to one plant per hill at the V5 stage to about 18 plants per plot. Natural foliar disease incidence and severity were evaluated at grain filling stage (R4-R5).

### Reaction to foliar diseases

The disease symptoms were identified using the CIMMYT field identification guide for maize diseases (CIMMYT, 2004). The diseases scored include NCLB, SCLB, CLS, MSV, SCR, GLS, and ALB. Disease symptoms were evaluated 3-4 weeks after silking by scoring affected areas of three leaves above and below ears (functional leaves for grain growth) of 10 plants in the central part of each plot using a 1 to 5 scale adapted from Badu-Apraku et al. (2012) and Wang et al. (2014): 1 = no lesions or scattered lesions, covering <5%; 2 = a few lesions on leaves, covering 6–10%; 3 = many lesions on leaves, covering 11–30%; 4 = large coalesced lesions covering 31–70%; and 5 = extensive, large, coalesced lesions covering almost the entire leaves, with many leaves dead. Based on the DS scores, the inbred lines were classified as R, MR, or S to each disease. Symptomatic leaves were sampled and transported to the laboratory for microbial analysis. Pathogens causing the observed diseases were isolated, identified, and confirmed based on morphological characters.

## Statistical analysis

The DS scores and the AUDPC data of the DLA were subjected to ANOVA using a fixed model [replicates and interactions among factors (inbred line × inoculation treatment)] in SAS/STAT v.9.4 (SAS Institute Inc., 2019). Means were separated with the Fisher’s protected least significant difference (LSD) test and standard errors were estimated for the interaction effects.

The field data (rating scale 1–5) were analyzed using a non-parametric ANOVA-type statistic (ATC) following the method of Shah & Madden (2004). Ratings were first converted to rank using Proc rank in SAS, and the ranked data were analyzed with mixed models (PROC MIXED/PROC GLIMMIX). Location, line, replication, and their interactions (Rep × Loc, Line × Loc) were considered as fixed effects, while replication (Loc) and block (Rep Loc) as random effects. Variance heterogeneities were examined and modeled (where appropriate) using the GROUP=Entry, following Stroup et al. (2018). Least significant difference (LSD) tests were performed on entry means, and contrast (High vs Low) were tested using lsmestimate.

## Results

### Detached Leaf Assay (DLA)

The PVA inbred lines varied in susceptibility to the evaluated pathogens in the DLA (Fig. 1). The progress of the lesions from the point of inoculation advanced slowly on leaves of certain inbred lines while in others the lesions appeared from day 2 onwards. Control leaves remained healthy throughout the experiments. When symptoms were present, lesion sizes caused by different isolates were similar at 4 DAI; however, some inbred lines showed no visible symptoms throughout the experiments. At 8 DAI, differences between R and S inbred lines were evident. High PVA inbred lines were in general more tolerant to CLS, SCLB, and NCLB, as per the DS score, RSI, and AUDPC values.

**Figure 1.**
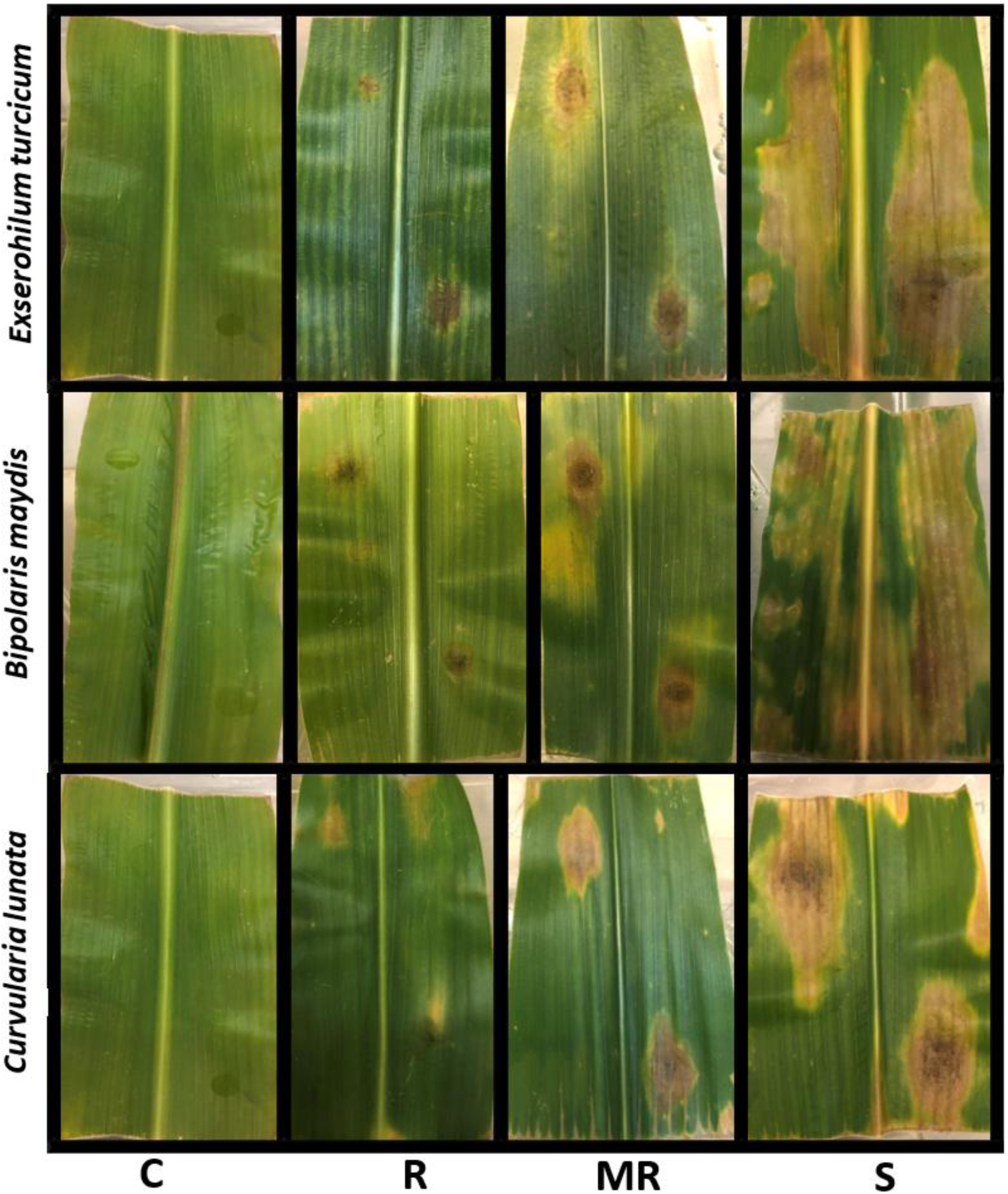
Representative reactions (R = resistant, MR = moderately resistant, S = susceptible, and C = non-inoculated control) of maize inbreds at 10 days after inoculation with three foliar pathogens, *Exserohilum turcicum, Bipolaris maydis,* and *Curvularia lunata*, using a detached leaf assay.

The ANOVA showed significant variation among the inbred lines for DS scores (*P* < 0.5–0.0001) and AUDPC values (*P* < 0.01–0.0001; Table 2). The inoculation treatment (pathogen) had a significant effect on DS scores and AUDPC (*P* < 0.0001). Further, the interactions of genotype × pathogen were significant for the measured traits (*P* < 0.05–0.01 and < 0.05–0.001, respectively) except for CLS. The contrast of the H vs L PVA revealed significant differences for DS scores (*P* < 0.01–0.0001) and AUDPC values (*P* < 0.05–0.001) except for CLS.

**Table 2.** Analysis of variance of disease severity (DS) scores and area under disease progress curve (AUDPC) values of PVA maize inbred lines inoculated with three foliar pathogens using a detached leaf assay.

| Source of variation | *Curvularia lunata*<sup>a</sup> |  |  | *Exserohilum turcicum*<sup>b</sup> |  | *Bipolaris maydis*<sup>c</sup> |  |  | Combined DS |  |  |
| --- | --- | --- | --- | --- | --- | --- | --- | --- | --- | --- | --- |
|  | DF | DS_10 | AUDPC | DS_10 | AUDPC | DF | DS_10 | AUDPC | DF | DS | AUDPC |
| Rep | 2 | 57.7† | 6,190.5† | 0.6 | 1.3 | 2 | 0.7 | 43.9† | 2 | 35.4† | 224.4† |
| Pathogen | 2 | 2,147.7† | 214.5† | 2,649.0† | 7,804.9† | 1 | 4,124.8† | 10,799.9† | 5 | 1,255.0† | 35,69.6† |
| Line | 22 | 1.5* | 7.4** | 1.6† | 6.5† | 22 | 0.8** | 3.4** | 22 | 3.7† | 15.0† |
| Line x Pathogen | 22 | 1.3 | 4.8 | 0.9* | 4.0*** | 22 | 0.8** | 3.4** | 110 | 1.10* | 4.5* |
| High vs. Low |  | -1.3 | -1.4 | -3.9† | -2.0* | 1 | -2.7** | -2.5* | 1 | -3.75*** | -2.7** |
| R<sup>2</sup> |  | 93.2 | 91.4 | 97.7 | 96.0 |  | 97.8 | 49.5 |  | 91.6 | 88.9 |
| CV |  | 21.5 | 24.6 | 12.1 | 16.0 |  | 16.4 | 13.5 |  | 15.2 | 18.0 |
| LSD |  | 0.6 | 0.4 | 0.9 | 0.9 |  | 0.5 | 1.1 |  | 0.4 | 0.8 |

\*, \*\*, \*\*\*, † significant at probability < 0.05, 0.01, 0.001, and 0.0001 levels, respectively. DS: Disease severity, AUDPC: Area under the disease progress curve; R<sup>2</sup>: coefficient of determination; CV: coefficient of variation; LSD: least significant difference.

At 10 DAI, the reaction of each PVA inbred line to *C. lunata* isolates were significant (*P* < 0.05) only for TZI1278, TZI2020, TZI2182, and TZI2116-3. The high PVA inbred lines TZI2116-3, TZI2182, TZI2065, and Tester 1 had the lowest DS scores than the non-PVA checks TZAR101 and TZAR102, while the low PVA inbred lines TZI1278, TZI2046 and Tester 2 had the highest DS scores than the other inbred lines. However, the high PVA inbred lines TZI2006 and TZI1653 had high DS scores for CLS. The same results were obtained with AUDPC values (with few changes in the order) because the DS scores and the AUDPC values were significantly correlated (r = 0.90, *P* < 0.0001; Table 3).

**Table 3.** Rank summation index (RSI) and resistance reaction of PVA maize inbreds inoculated with various isolates of three foliar pathogens.

| Inbreds | PVA | *Curvilaria lunata*<sup>a</sup> |  |  | *Exserohilum turcicum*<sup>b</sup> |  |  | *Bipolaris maydis*<sup>c</sup> | RSI<sup>d</sup> | Class<sup>e</sup> |
| --- | --- | --- | --- | --- | --- | --- | --- | --- | --- | --- |
|  |  | CV1 | CV2 | Average | ET1613 | ET1612 | Average | SBL05 |  |  |
| IITATZI2071 | 43.3 | 9.7 | 13.1 | 11.4 ebdacf | 12.4 | 12.8 | 12.6 ejhigf | 12.1 bdec | 155 | R |
| IITATZI1715 | 45.8 | 11.3 | 13.4 | 12.3 bac | 12.5 | 13.3 | 12.9 edhigcf | 12.7 bdac | 209 | MR |
| IITATZI2066 | 43.4 | 12.0 | 11.6 | 11.8 bdac | 12.4 | 12.3 | 12.4 jhig | 12.5 bdac | 170 | MR |
| IITATZI2065 | 36.5 | 10.6 | 11.2 | 10.9 ebdcf | 13.3 | 12.2 | 12.8 ejdhigf | 11.6 bdec | 149 | R |
| IITATZI2163 | 27.8 | 12.0 | 12.4 | 12.2 bac | 13.7 | 12.9 | 13.3 ebdhagcf | 12.4 bdac | 196 | MR |
| IITATZI2142-1 | 35.7 | 11.2 | 10.3 | 10.7 edcf | 12.0 | 12.0 | 12.0 jhi | 10.9 dec | 125 | R |
| IITATZI1653 | 51.7 | 11.2 | 12.5 | 11.8 bdac | 12.8 | 13.6 | 13.2 ebdhgcf | 12.3 bdac | 206 | MR |
| IITATZI2182 | 37.0 | 9.8 | 9.6 | 9.7 f | 12.7 | 12.3 | 12.5 jhigf | 11.9 bdec | 142 | R |
| IITATZI2006 | 31.4 | 11.4 | 13.3 | 12.4 bac | 12.8 | 14.4 | 13.6 ebdagcf | 12.4 bdac | 213 | MR |
| IITATZI1278 | 13.4 | 12.3 | 13.1 | 12.7 ba | 13.6 | 12.1 | 12.8 edhigf | 14.3 a | 233 | S |
| IITATZI2068 | 5.4 | 11.0 | 12.3 | 11.6 ebdac | 13.8 | 14.0 | 13.9 ebdac | 12.6 bdac | 196 | MR |
| IITATZI2116-3 | 36.3 | 10.2 | 9.3 | 9.8 ef | 11.6 | 12.5 | 12.0 jhi | 10.6 de | 113 | R |
| IITATZI2019 | 14.6 | 12.6 | 11.9 | 12.2 bac | 14.9 | 13.6 | 14.2 bac | 13.3 ba | 254 | S |
| IITATZI2005-1 | 27.0 | 11.6 | 12.2 | 11.9 bdac | 14.5 | 14.3 | 14.4 ba | 12.8 bac | 230 | S |
| IITATZI2046 | 19.0 | 12.3 | 12.6 | 12.4 bac | 15.0 | 14.3 | 14.7 a | 13.5 ba | 265 | S |
| IITATZI2028 | 15.8 | 12.2 | 12.7 | 12.5 bac | 14.0 | 14.0 | 13.9 bdac | 14.3 a | 265 | S |
| IITATZI2012-2 | 22.3 | 12.3 | 12.3 | 12.3 bac | 10.9 | 12.6 | 11.8 ji | 10.1 e | 153 | R |
| IITATZI2026-3 | 12.8 | 12.8 | 9.4 | 11.1 ebdacf | 13.6 | 11.0 | 12.3 jhig | 12.9 bac | 186 | MR |
| IITATZI2020 | 8.7 | 12.6 | 13.0 | 12.8 a | 13.6 | 13.6 | 13.6 ebdagcf | 13.3 ba | 251 | S |
| TZAR101 | 0.0 | 9.5 | 10.7 | 10.1 edf | 13.6 | 12.6 | 13.1 ebdhgcf | 13.4 ba | 165 | MR |
| TZAR102 | 0.0 | 9.7 | 12.8 | 11.2 ebdacf | 15.2 | 12.2 | 13.7 ebdacf | 13.6 ba | 175 | MR |
| Tester 1 | 14.0 | 9.9 | 10.5 | 10.2 edf | 12.7 | 11.4 | 12.0 jhi | 12.0 bdec | 116 | R |
| Tester 2 | 25.0 | 12.5 | 12.6 | 12.5 bac | 12.8 | 10.1 | 11.5 j | 12.0 bdec | 175 | MR |
| Mean |  | 11.3 | 11.9 | 11.6 | 13.2 | 12.8 | 13.0 | 12.5 | 189 |  |
| SE<sup>f</sup> |  | 4.6 | 4.8 | 4.1 | 4.1 | 3.7 | 4.5 | 4.1 |  |  |
| CV (%)<sup>g</sup> |  | 17.7 | 20.8 | 18.1 | 14.4 | 13.5 | 13.0 | 14.7 |  |  |
| LSD<sup>h</sup> |  |  |  | 1.69 |  |  | 1.36 | 1.93 |  |  |

<sup>a</sup> AUDPC values of the two evaluated isolates (CV1 and CV2) at 10 days after inoculation (DAI).
<sup>b</sup> AUDPC values of the two evaluated isolates (NGET16-IK-12 and NGETIB-16-13) at 10 DAI.
<sup>c</sup> AUDPC values of the *B. maydis* isolates evaluated (SLB-05) at 10 DAI.
<sup>d</sup> RSI: rank summation index.
<sup>e</sup> Overall reaction to the three foliar pathogens. R: resistant; MR: moderately resistant; S: susceptible
<sup>f</sup> SE: standard error.
<sup>g</sup> CV%: Coefficient of variation expressed in percentage.
<sup>h</sup> Least significant difference.

The inbred lines reacted relatively similarly to the two *C. lunata* isolates with no significant difference recorded for DS and AUDPC of CV1 and CV2. For NCLB, the reactions of each inbred line to both isolates were similar (*P* > 0.05) except for Tester 2, TZI2012-2, TZI2005-1, and TZI2046. Tester 2 recorded the lower AUDPC value for *E. turcicum* when inoculated with NGETIK16-12, followed by TZI2012-2, TZI2142-1, and TZI2116-3. However, Tester 2 inoculated with the virulent isolate of *E. turcicum* NGETIB16-13 had the highest AUDPC value (Table 3). Four of the five inbred lines (TZI2068, TZI2028, TZI2019, and TZI2046) with the highest AUDPC values are among the lowest PVA inbred lines. Complementing this group is TZI2005-1, a high PVA inbred line. The AUPDC values of these five inbred lines were higher when inoculated with NGETIB16-13 indicating higher virulence of NGETIB16-13 over NGETIK16-12. On the other hand, TZI2012-2, TZI2116-3, and TZI2006 were more susceptible to NGETIK16-12 than to NGETIB16-13 (Table 3). The reactions of each inbred line to *B. maydis* isolates were not significantly different (*P* > 0.05) except for TZI2012-2, TZI2116-3, TZI1278, and TZI2028. The inbred lines TZI2012-2, TZI2116-3, TZI2142-1, TZI2065, TZI2182, and Tester 1 had the lowest DS score and AUDPC values compared to the control TZAR101 and TZAR102. They are all high PVA inbred lines except TZI2012-2. Further, the low PVA inbred lines TZI2046, TZI1278, TZI2028, TZI2026-3, TZI2020, and TZI2019, and the non-PVA TZAR101 and TZAR102 had the highest DS score and AUDPC value, indicating the tolerance of high PVA to *B. maydis* infection.

The RSI values revealed some inbred lines having high levels of resistance to one or more pathogens, others were MR, and some were S to one or more pathogens. None of the inbred lines recorded an RSI value below 100. The RSI ranged from 113 to 265. Inbred lines were classified in three groups as follows: R (RSI range = 113–155), MR (RSI range = 165–213), and S (RSI range = 230–265) to all foliar pathogens (Table 3). In all, 30.4% of the inbred lines were classified as R, 43.5% as MR, and 26.1% as S to all foliar pathogens. Inbred lines with the lowest RSI classified as R are high PVA inbred lines, except TZI2012-2. All the inbred lines classified as S have low PVA, except TZI2005-1. Additionally, the high PVA inbreds TZI2066, TZI1653, and TZI1715 (43 to 51 µg/g) are classified as MR. The high PVA inbreds TZI2116-3, TZI2142-1, TZI2182, and TZI2065 had the lowest DS score and AUDPC value for the three pathogens (Table 3).

### Field assessment

Multiple foliar diseases were observed and scored in the Ikenne and Ibadan fields. Pathogens such as *P. polysora*, *B. maydis*, *C. lunata*, *C. zeae-maydis*, and *Colletotrichum* spp. were identified from the samples collected from the field and scored as SCR, SCLB, CLS, GLS, and ALB diseases, respectively.

The ANOVA of the DS scores gave similar trends in the two locations (except for GLS) although the infection level was higher at Ikenne for most diseases (Table 4). The levels of significance for sources of variation for the traits were similar for both locations allowing the analysis of the combined data. The mean DS scores of the foliar diseases of the PVA inbred lines varied significantly at each location and across location (*P* < 0.5–0.0001). In the combined analysis, location was significant for CLS, GLS, and ALB, while line × location interaction was not significant. The contrast of H vs L PVA revealed a significant difference for DS scores. The contrast H vs L was negative for MSV (*P* < 0.0001), CLS, GLS, and ALB (*P* < 0.01) indicating that high PVA inbred lines had less susceptibility to these diseases than the low PVA inbred lines. However, for SCR (*P* < 0.0001) and SCLB, the contrast was positive. These results were similar within and across locations (Table 4).

**Table 4.**
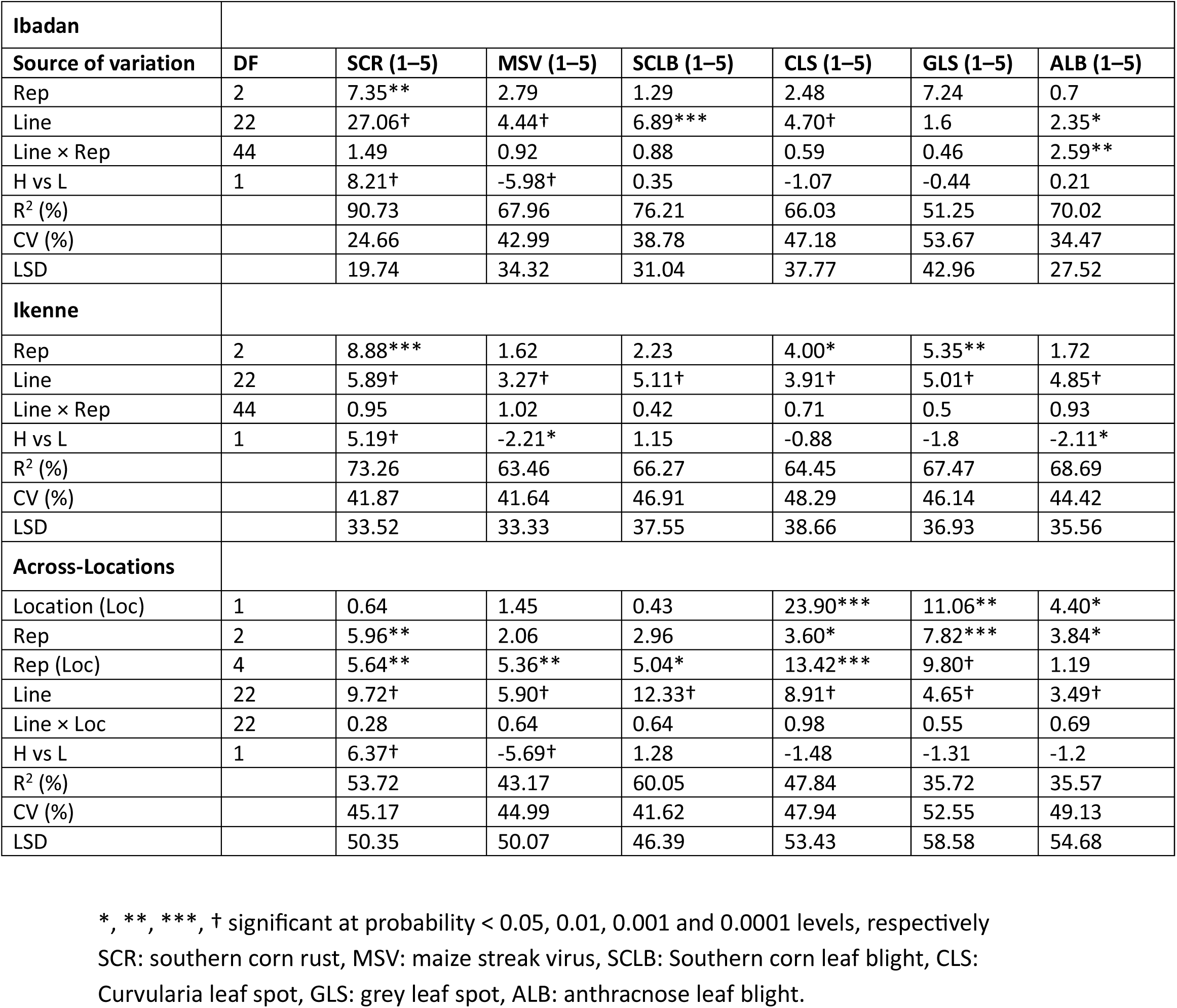
Analysis of variance of the disease severity scores in PVA maize inbred lines tested in field conditions at Ikenne and Ibadan, Nigeria.

| **Ibadan** |  |  |  |  |  |  |  |
| --- | --- | --- | --- | --- | --- | --- | --- |
| **Source of variation** | **DF** | **SCR (1–5)** | **MSV (1–5)** | **SCLB (1–5)** | **CLS (1–5)** | **GLS (1–5)** | **ALB (1–5)** |
| Rep | 2 | 7.35** | 2.79 | 1.29 | 2.48 | 7.24 | 0.7 |
| Line | 22 | 27.06† | 4.44† | 6.89*** | 4.70† | 1.6 | 2.35* |
| Line × Rep | 44 | 1.49 | 0.92 | 0.88 | 0.59 | 0.46 | 2.59** |
| H vs L | 1 | 8.21† | -5.98† | 0.35 | -1.07 | -0.44 | 0.21 |
| R<sup>2</sup> (%) |  | 90.73 | 67.96 | 76.21 | 66.03 | 51.25 | 70.02 |
| CV (%) |  | 24.66 | 42.99 | 38.78 | 47.18 | 53.67 | 34.47 |
| LSD |  | 19.74 | 34.32 | 31.04 | 37.77 | 42.96 | 27.52 |
| **Ikenne** |  |  |  |  |  |  |  |
| Rep | 2 | 8.88*** | 1.62 | 2.23 | 4.00* | 5.35** | 1.72 |
| Line | 22 | 5.89† | 3.27† | 5.11† | 3.91† | 5.01† | 4.85† |
| Line × Rep | 44 | 0.95 | 1.02 | 0.42 | 0.71 | 0.5 | 0.93 |
| H vs L | 1 | 5.19† | -2.21* | 1.15 | -0.88 | -1.8 | -2.11* |
| R<sup>2</sup> (%) |  | 73.26 | 63.46 | 66.27 | 64.45 | 67.47 | 68.69 |
| CV (%) |  | 41.87 | 41.64 | 46.91 | 48.29 | 46.14 | 44.42 |
| LSD |  | 33.52 | 33.33 | 37.55 | 38.66 | 36.93 | 35.56 |
| **Across-Locations** |  |  |  |  |  |  |  |
| Location (Loc) | 1 | 0.64 | 1.45 | 0.43 | 23.90*** | 11.06** | 4.40* |
| Rep | 2 | 5.96** | 2.06 | 2.96 | 3.60* | 7.82*** | 3.84* |
| Rep (Loc) | 4 | 5.64** | 5.36** | 5.04* | 13.42*** | 9.80† | 1.19 |
| Line | 22 | 9.72† | 5.90† | 12.33† | 8.91† | 4.65† | 3.49† |
| Line × Loc | 22 | 0.28 | 0.64 | 0.64 | 0.98 | 0.55 | 0.69 |
| H vs L | 1 | 6.37† | -5.69† | 1.28 | -1.48 | -1.31 | -1.2 |
| R<sup>2</sup> (%) |  | 53.72 | 43.17 | 60.05 | 47.84 | 35.72 | 35.57 |
| CV (%) |  | 45.17 | 44.99 | 41.62 | 47.94 | 52.55 | 49.13 |
| LSD |  | 50.35 | 50.07 | 46.39 | 53.43 | 58.58 | 54.68 |

\*, \*\*, \*\*\*, † significant at probability < 0.05, 0.01, 0.001 and 0.0001 levels, respectively SCR: southern corn rust, MSV: maize streak virus, SCLB: Southern corn leaf blight, CLS: Curvularia leaf spot, GLS: grey leaf spot, ALB: anthracnose leaf blight.

### Frequency of the foliar diseases and inbred lines reaction

The frequency of the multiple foliar diseases within and across location is presented in Fig. 2. SCR was the most predominant with infection scored in 30.3%, 21.7%, and 47.8% at Ibadan, Ikenne, and across locations, respectively. The frequency of MSV was 13.0% and 4.4%, SCLB 17.4% and 13.0%, CLS 8.7% and 13.0%, GLS 8.7% and 17.4%, and ALB 4.3% and 4.3% at Ibadan and Ikenne, respectively. No symptoms of NCLB were detected in the fields.

**Figure 2.**
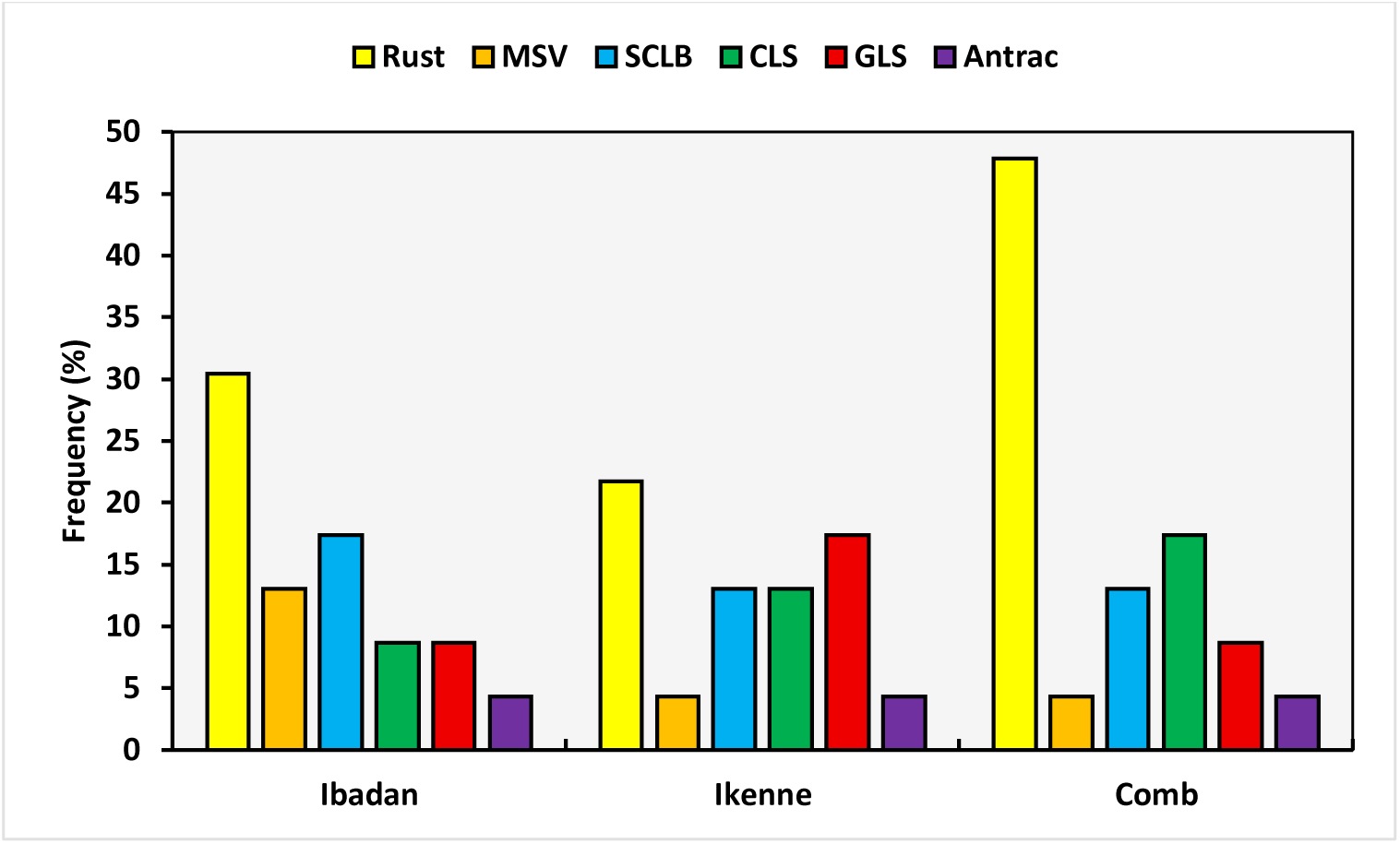
Frequencies of naturally occurring foliar diseases within and across locations affecting the PVA maize inbred lines.

The proportions of lines that were R, MR, or S to natural disease infection is presented in Table 5. The DS scores varied significantly among the inbred lines (Table 4). However, the inbred lines were generally R to MR to the foliar diseases except to SCR. The DS scores ranged from 1.00 to 2.28 for MSV, 1.08 to 2.94 for SCLB, 1.09 to 2.90 for GLS, 1.00 to 1.52 for ALB, and 1.28 to 4.16 for CLS (Table 5). The lines TZI2065, TZI2026-3, and TZI1278 were classified S to CLS with DS scores 3.97, 3.73 and 4.16, respectively.

**Table 5.**
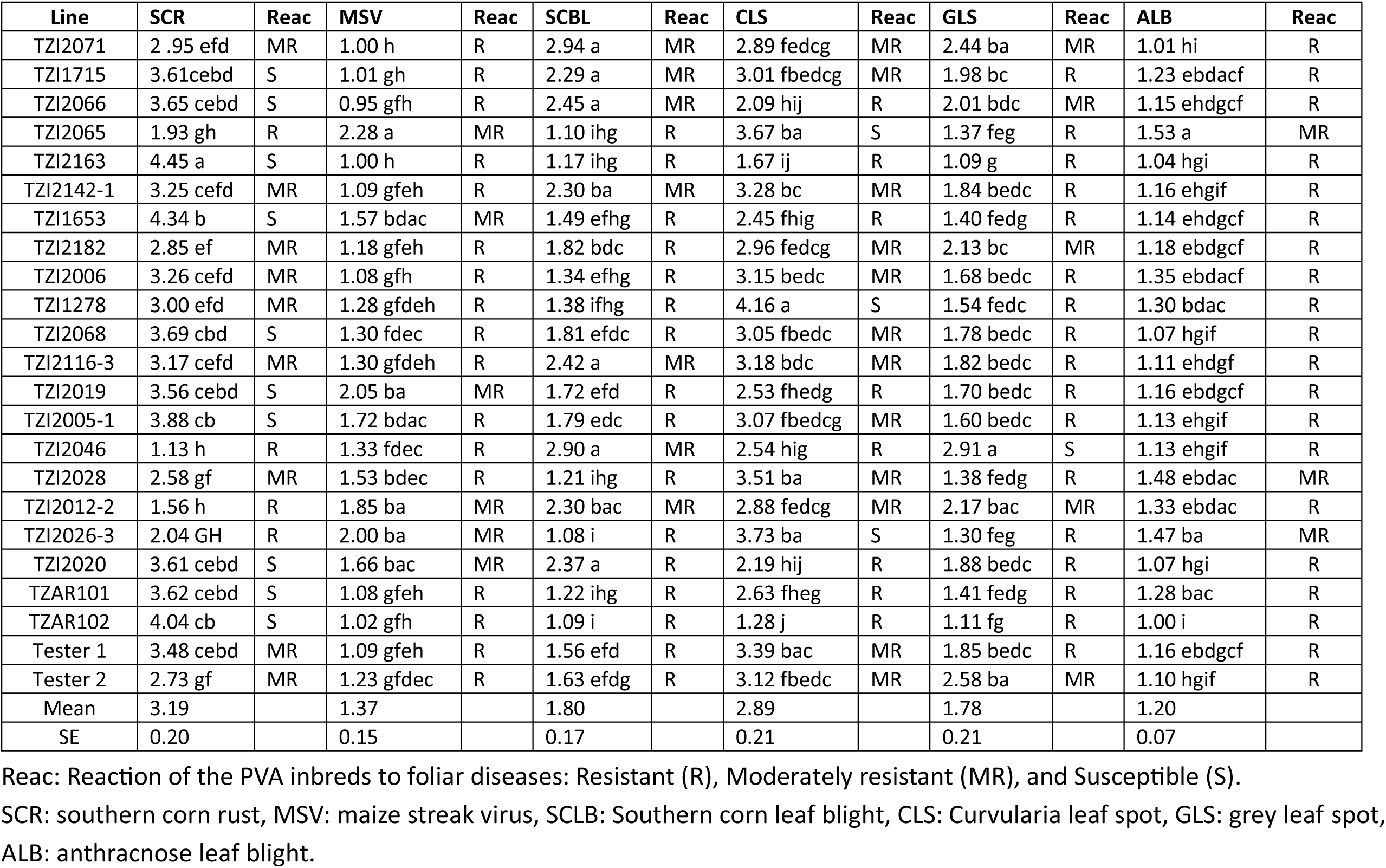
Mean disease scores and reaction of PVA maize inbred lines to multiple foliar diseases.

Southern corn rust (SCR) was the most severe among the foliar diseases with 43.5% of the inbred lines classified as S and DS scores ranging from 1.12 to 4.45. Overall, 17.4% were R and 39.1% were MR to SCR. TZI2046, TZI2012-2, TZI2065, and TZI2026-3 were the most resistant to SCR while TZI1653 and TZI2163 were the most susceptible with a DS score of 4.34 and 4.45, respectively. The check inbreds TZAR102 and TZAR101 were classified as S to SCR while R to the other diseases.

The percentage of high and low PVA inbred lines infected by the pathogens ranged from 8.3% to 41.7% and 12.2% to 37.5%, respectively. The high PVA inbred lines were more resistant to MSV, CLS, and ABL than the low PVA inbred lines while the later were less susceptible to SCR, SCLB, and GLS (Fig. 3). Over 50% of the lines were R to MR to multiple foliar diseases with a DS score ranging from 1.00 to 3.48. The four R lines to SCR, the most virulent and spread in the field, were also R to MR to the other diseases except TZI2065 and TZI2026-3 which were S to CLS (Table 5).

**Figure 3.**
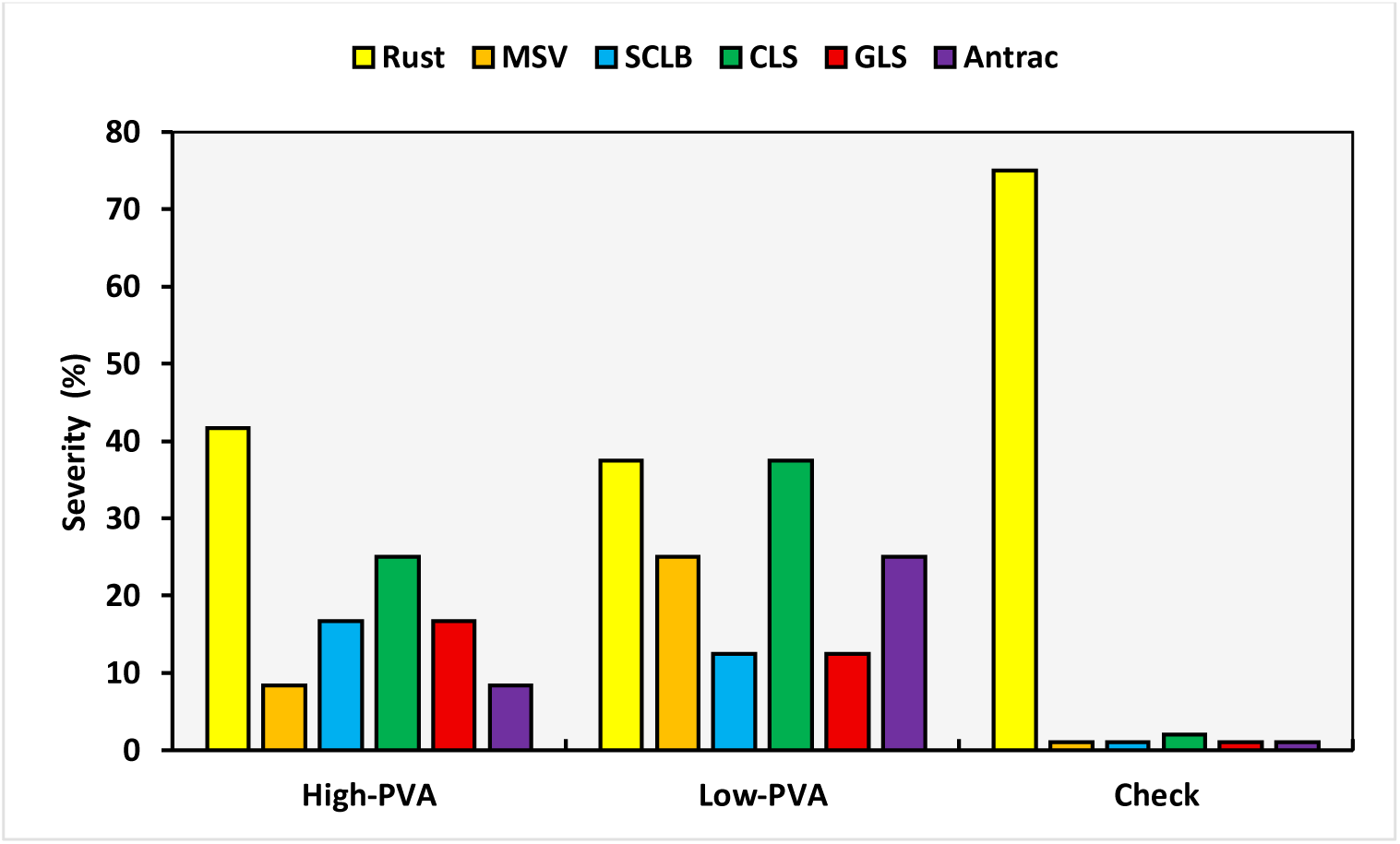
Disease severity percentage of the high and low PVA maize inbred lines in response to foliar pathogens under natural conditions.

## Discussion

In the current study, we evaluated maize PVA inbred lines in DLA for their resistance to *E. turcicum*, *B. maydis*, and *C. lunata*, pathogens causing damaging foliar diseases. In addition, the inbred lines were evaluated in field trials for resistance to multiple foliar diseases under natural infection. The PVA maize inbred lines were classified as R, MR, and S to the diseases based on an RSI calculated with AUDPC values obtained from the 1^st^ to 5^th^ time DS scores. Seven inbred lines were classified as R, 10 as MR, and six as S to all foliar diseases. The high PVA inbred lines TZI2116- 3, TZI2142-1, TZI2182, and TZI2065 had good levels of resistance in DLA to all foliar diseases. Under natural field infestation, SCR was the most prevalent disease, with 43.5% of the inbred lines classified as S. High PVA inbred lines showed greater resistance to MSV, CLS, and ABL, while low PVA inbred lines were less affected by SCR, SCLB, and GLS. Over 50% of the evaluated inbred lines were R to MR to multiple foliar diseases.

Screening for multiple disease resistance, including natural infection in hot spots and artificial inoculation has consistently been part of efforts of IITA’s MIP to develop improved maize cultivars with utility in SSA. Sometimes these efforts are constrained because certain diseases are specific to particular agroecological zones (Badu-Apraku & Fakorede, 2017). Additionally, climate change has increased the distribution, frequency, and severity, resulting in disease outbreaks, especially GLS that was confined to the mid-altitudes of Plateau State (Menkir & Ayodele, 2005).

Detached-leaf assay’s (DLA) have been used to screen maize and other crops for resistance to foliar pathogens, facilitating the identification of genotypes resistant to multiple diseases (Degani & Cernica, 2014; Alakonya et al., 2018; Aregbesola et al., 2020; Bankole et al., 2022). They have generally produced similar results to those obtained from screenhouse and field tests allowing identification of both highly susceptible and highly resistant cultivars (Bouhassan et al., 2004; Aregbesola et al., 2020). Therefore, the DLA can serve as a useful tool for initially screening large number of genotypes to effectively identify resistant genotypes. This also reduces the costs associated with screenhouse or field tests and significantly minimizes the risks of obtaining ambiguous results due to fluctuating environmental conditions, co-infection of diverse pathogens, and insect damage. However, according to Bouhassan et al. (2004), the DLA method partially suppresses the plant’s intrinsic physiological reactions and its interaction with the environment by removing the leaves, making it necessary to confirm results in field tests.

The significant pathogen and line effects, and line × pathogen interaction observed for both SCLB and NCLB DS score indicates that i) inbred lines exhibit differential responses to foliar pathogen infection and ii) varying levels of virulence exist among the foliar pathogens. This was shown by the inbred lines displaying higher AUDPC value when inoculated with NGETIB16-13 compared to NGETIB16-12. Some inbred lines were resistant to one isolate of a pathogen and susceptible to the other while other inbred lines were resistant to both isolates (Table 3). These results are consistent with those reported in other studies (Aregbesola et al., 2020; Bankole et al., 2022). Therefore, the advantage of using different isolates of a pathogen or a mixture of isolates resides in identifying broad resistance to foliar pathogens. However, according to Bouhassan et al. (2004), the use of a single virulent isolate inoculum, rather than a mixture of isolates with varying pathogenicity, can avoid confusion between vertical and horizontal resistance.

The RSI ranks the inbred lines for their reaction to the foliar diseases. However, no inbred lines recorded RSI values below 100 indicating that none was completely resistant or highly resistant to all foliar pathogens. The inbred lines classified as R to all foliar pathogen used in the current study contain high PVA while those classified S contain low PVA, except inbred TZI2005-1 (Table 3). However, the resistance observed in the low PVA inbred lines such as TZI2012-2, as indicated by its low RSI, could be attributed to factors such as leaf architecture defense, inducible defense responses (e.g., reactive oxygen species), a unique genetic background, or the presence of specific resistance QTL unrelated to carotenoid biosynthesis. The contrast H vs L revealed a significant difference between the high PVA and low PVA inbred lines for DS scores and AUDPC values indicating that PVA maize are more resistant to foliar diseases compared to low PVA and non-PVA maize. The AUDPC values of the DLA experiments clearly confirmed the reaction (R to MR) of most of the inbred lines based on the DS score in the field.

Detecting high PVA inbred lines resistant to all tested foliar pathogens suggest that these inbreds possess QTLs conferring resistance to multiple diseases (Martins et al., 2019). Indeed, the chromosomal region bin 4.08 was reported containing a QTL associated with resistance to AFL and contains a cluster of resistant genes against multiple diseases (Wisser et al., 2006; Martins et al., 2019) such as NCLB (Balint-Kurti et al., 2010), and maize rough dwarf disease (Wang et al., 2019). For those superior inbred lines, it should be further investigated the genetic basis conferring resistance to *A. flavus* colonization, AFL accumulation, and to multiple foliar diseases.

The significant genotypic effect for the DS scores within and across location indicated variation in reaction to different foliar diseases in the PVA maize inbred lines. The lines classified as R to MR had DS score of 1 and 2 (R), and 3 (MR), respectively, while those S had rating score of 4. No inbred line was R to all the foliar diseases. However, the majority of them were R to MR to multiple diseases.

The average DS scores for all the diseases were relatively low (ranging from 1.4 to 2.9), except for SCR (3.19) indicating that the PVA inbred lines have tolerance to multiple foliar diseases. The lowest disease and the predominance of SCR results concur with those reported by Iseghohi et al. (2024) who evaluated PVA-enriched maize genotypes for foliar diseases tolerance, and by Akinwale and Oyelakin (2018) on early and extra-early maturing tropical maize inbred lines under natural infection field trials. The low DS scores for MSV, SCLB, GLS, and ALB indicates that these PVA inbred lines have been enhanced for resistance to multiple diseases and as stated earlier (Badu-Apraku & Fakorede, 2017), confirming the importance of routine diseases resistance screening in breeding program. However, the maximum DS scores for CLS and SCR exceeded 3.0 and 4.0, respectively, indicating susceptibility in certain PVA inbred lines to these diseases. These results align with those of Akinwale and Oyelakin (2018), who reported SCR and CLS exceeding the susceptibility threshold, with the highest value for disease progression (*P* = 0.52 and 0.27, respectively). In addition, the incidence of SCR was higher both within and across locations, as a projection study of climate change effects revealed that certain regions of Africa will likely experience an increased risk of occurrence of both common rust and SCR (Ramirez-Cabral et al., 2017), and urge to take proactive measures to mitigate the potential risks. According to Iseghohi et al. (2024), MSV and SCR can reduce genotypic variability for lutein by 36.7% and 18.7%, respectively, although they do not significantly affect PVA content. Hence, the identified resistant lines to SCR can be used as donor parents to improve PVA maize resistance to SCR and other foliar diseases. Additionally, several of these inbred lines such as TZI2071, TZI1715, TZI1653, and TZI2005-1, which are classified as R to MR to multiple foliar diseases, were also found to be resistant to both Aspergillus and Fusarium ear rots and tolerant to aflatoxin and fumonisin accumulation (Mboup et al., 2024). Inbreds TZI1715, TZI1653, and TZI2071 combine desirable negative GCA effect for *A. flavus* infection and aflatoxin accumulation and significant positive GCA effect for beta carotene (BC) and PVA content (Mboup et al., 2023). Therefore, these inbred lines are promising candidates for developing high PVA hybrids that are tolerant to foliar diseases and reduce mycotoxin exposure to be tested in multi-location trials for increase production, contributing to global food and nutrition security and sustainability.

The AUDPC values observed in the PVA inbred lines, when inoculated with *C. lunata* and *B. maydis* in the DLA, clearly confirmed the R to MR of most of the inbred lines, as determined with the field DS score. However, TZI2065 and TZI2026-3 classified as S to CLS in the field, were among the inbred lines ranked resistant to C. lunata, while lines that were S to *B. maydis* in the DLA were R to MR to SCLB in the field. These differences could be due to fluctuating climatic conditions influencing disease development, spread of the pathogen in the field, and virulence levels of the isolates used during the DLA.

The contrast H vs L was significant only for MSV (-4.65, *P* < 0.0001), with high PVA inbred lines allowing less infection than the low PVA, and for SCR (9.36, *P* < 0.0001) where the low PVA inbred lines were less susceptible. However, in the DLA, the contrast H vs L was significant for DS scores and AUDPC values of all the fungi except for *C. lunata*. In addition, as the effects of foliar diseases on grain carotenoids in maize have not been established, a large-scale trial involving artificial inoculation of each disease pathogen in prevalent agroecological zones is necessary to ascertain and establish the resistance levels of PVA inbred lines to multiple diseases, to determine the effect of carotenoids content on the diseases, and facilitate the selection of PVA inbred lines for breeding PVA maize with higher levels of foliar disease resistance.

## Conclusion

Several foliar diseases threaten maize production across diverse agroecological zones within SSA. The current study evaluated PVA maize inbred lines for resistance to several foliar diseases using both DLA and natural field infestation. Several inbred lines were R to certain diseases while being MR or S to others, with high PVA inbreds TZI2116-3, TZI2142-1, TZI2182, and TZI2065 exhibiting the lowest DS ratings and AUDPC values for *C. lunata*, *B. maydis*, and *E. turcicum*. The contrast H vs L PVA revealed that high PVA maize possess higher levels of resistance to foliar diseases compared to low PVA and non-PVA maize. The DLA effectively discriminated resistant and susceptible inbred lines, demonstrating its potential as reliable tool for large-scale screening of genotypes in breeding programs, thereby reducing the time and cost associated with field trials. However, further research should explore large number of PVA maize inbred lines collection and validate the resistance of selected genotypes through artificial inoculation in disease hotspots where each pathogen is predominant.

## Conflict of interest

The authors declare that the research was conducted in the absence of any commercial or financial relationships that could be construed as a potential conflict of interest.

## Funding

This work is part of a PhD project of the first author, funded by the African Union Commission through the Pan African University, and One CGIAR Plant Health Initiative supported by the CGIAR Trust Fund contributors (https://www.cgiar.org/funders/).

## CRediT authorship contribution statement

Conceptualization: MM, AO-B, AM; Methodology: MM, ID, AM, AO-B; Supervision: AOM-M, BO, AM, AO-B; Data analysis: MM, ID; Manuscript draft: MM, AM, AO-B; Resources: MM, AM, AO-B; Manuscript review and editing: MM, AOM-M, BO, WM, SM, ID, AM, AO-B.

**M. Mboup:** Writing – review & editing, Writing – original draft, Visualization, Validation, Methodology, Investigation, Funding acquisition, Formal analysis, Data curation, Conceptualization. **A.O. Aduramigba-Modupe:** Writing – review & editing, Supervision, Resources. **B. Olasanmi:** Writing – review & editing, Supervision, Resources, Investigation. **W. Mengesha:** Writing – review & editing, Resources, Methodology, Investigation. **S. Meseka:** Writing – review & editing, Resources, Methodology. **I. Dieng:** Writing – review & editing, Visualization, Validation, Software, Resources, Methodology, Formal analysis, Data curation. **A. Menkir:** Writing – review & editing, Supervision, Resources, Project administration, Methodology, Funding acquisition, Conceptualization. **A. Ortega-Beltran:** Writing – review & editing, Writing – original draft, Supervision, Resources, Project administration, Methodology, Investigation, Funding acquisition, Conceptualization.

## Acknowledgements

The authors appreciate the staff of Maize Improvement Program at IITA for their technical assistance in the field research, and staff of IITA Pathology Laboratory and Mycotoxin Unit for their contributions to the execution of this research.

## Abbreviations

ALB: anthracnose leaf blight
AUDPC: area under the disease progress curve
CLS: Curvularia leaf spot
CIMMYT: International Maize and Wheat Improvement Center
DLA: detached leaf assay
DS: disease severity
GLS: grey leaf spot
H: High
PVA: Consider HPVA
IITA: International Institute of Tropical Agriculture
L: Low
PVA: Consider LPVA
MIP: maize improvement program
MR: moderately resistant
MSV: maize streak virus
NCLB: northern corn leaf blight
PVA: provitamin A
QTL: quantitative trait loci
R: resistant
RSI: rank summation index
S: susceptible
SCLB: Southern corn leaf blight
SCR: southern corn rust
VAD: Vitamin A deficiency

## Notes

### Competing Interest Statement

The authors have declared no competing interest.

